# Automatic Integration of Gender Information during Phrase Processing: ERP Evidence

**DOI:** 10.1101/2020.07.16.206045

**Authors:** Maria Alekseeva, Andriy Myachykov, Beatriz Bermudez Margaretto, Yury Shtyrov

## Abstract

Both linguistic (e.g., words, syntax) and extralinguistic (e.g., voice quality) information needs to be considered by interlocutors during linguistic communication. The effects of extralinguistic information on neural sentence processing are particularly poorly understood. Here, we used EEG and passive non-attend design with visual distraction in order to investigate how extralinguistic information affects brain activity during syntactic processing. We collected ERPs while participants listened to Russian pronoun-verb phrases recorded in either male or female voice. We manipulated congruency between the grammatical gender signaled by the verb’s ending and the speaker’s apparent gender. We registered both early and late phrase processing signatures in the incongruent conditions including ELAN (peaking at ∼150 ms) and N400. Our data suggest a high degree of automaticity in integrating extralinguistic information during syntactic processing indicating existence of a rapid automatic syntactic integration mechanism sensitive to both linguistic and extralinguistic information.

## Introduction

During linguistic communication, we do not only rely on linguistic information (phonology, lexical semantics, grammar, etc.) but also on extralinguistic (or pragmatic) information provided by the speaker and the conversation context (see [1-2] for a review). Some of this information can be extracted from the speaker’s voice, and it may be particularly relevant for interpreting the message as it projects speaker-related characteristics including identity, age, gender, and emotional state. Existing evidence suggests that this information can directly modulate sentence processing. For example, listener’s expectations about the speaker’s language can help to decode the speech signal and predict upcoming information [3-4]. Yet, although many studies examined different psycho- and neurolinguistics processes underpinning language comprehension, little is known about the involvement of speaker-related features during this process.

The speaker’s voice provides the listener with a quick and reliable source of information that can facilitate sentence comprehension [5]. Among the first key characteristics, which can be identified from the speaker’s voice is their *apparent gender* [6-11]. Junger et al. [8] registered voice gender processing correlates in cingulate cortex and bilateral inferior frontal gyri with increased activation for opposite sex in fronto-temporal regions. The latter suggests that in order to make sense of the message in addition to evaluating the speaker’s gender, the listener needs to consider their own gender. Finally, many languages mark the *grammatical gender* explicitly adding another source of gender information and further complicating the interplay between these factors. Attempts to investigate how the listener integrates gender information from these different sources have shown, for example, that listeners process sentences faster when the grammatical gender is congruent with their own gender [12] resulting in reaction time decrease in congruent conditions (especially for females). An ERP study of semantically gendered words also showed an enhanced mismatch negativity (MMN) response to words whose semantic gender matched the gender of the speaker [13], presumably reflecting facilitated activation of the underlying memory traces. Furthermore, an interaction between the speaker’s and the listener’s genders during the sentence comprehension was documented as an early (at around 150 – 250 ms) increase of the MMN amplitude in listeners presented with opposite gender voices, compared the gender-matching presentations [13].

Other ERP studies have reported neurophysiological responses to inconsistencies between the message meaning and the speaker’s representation, typically manifested as a modulation of the N400 and/or P600 components [14-16]. Diverging patterns of ERP results found in these studies are likely related to the nature of the used mismatch manipulations; whereas the P600 component ─ typically associated with syntactic incongruences ─ is found under violations of stereotypical role nouns (e.g., *face powder, fight club* by male and female voices, respectively), the semantically-related N400 effect is found under semantic-pragmatic incongruences (e.g., “*I am going to the night club”* by child’s voice).

Understanding how gender information is integrated by the listeners is particularly important when one considers the differences in how different languages signal grammatical gender. In some languages, such as in English or Mandarin, overt gender marking is almost completely lacking. Many other languages including most Slavic languages explicitly mark grammatical gender in nouns, verbs, and adjectives, often in a complicated interdependent manner. Russian is one of such languages, offering an optimal testbed for investigating linguistic and extralinguistic gender integration. As far as we know, there is only one study addressing this question in a Slavic language [17]. Using Slovak, Hanulíková & Carreiras [17] found that, during an active-listening task, the integration of speaker-related information and morphosyntactic information occurred late during complex sentence processing. Additionally, a conflict between the speaker’s and the word’s genders (e.g., ‘*I ∗stole*_*MASC*_ *plums’* in female voice) was reflected in the modulation of the N400 component. Given that N400 modulations have been consistently found for morphosyntactic violations, in particular, number [18-19], person [20], and gender agreement [21] as well as in phrase structure violations [22], this result may suggest that extralinguistic information is directly integrated during online (morpho)syntactic processing.

However, N400 is also known to be related to conscious top-down controlled integration of linguistic information [23-24]. Indeed, in the study described above, the participant’s overt attention to the stimuli was required, and the effect appeared rather late in the comprehension processes. Thus, the question still remains whether such findings reflect the involvement of genuine online parsing mechanisms or secondary post-comprehension processes (such as repair and reanalysis, [23, 25]. Importantly, syntactic parsing was shown to commence much earlier and to take place in a largely automatic fashion, as demonstrated in studies using early left-anterior negativity (ELAN) or syntactic MMN. In particular, ELAN modulation around 200 ms has been reported during outright violations of the obligatory structure reflecting an automatic early analysis of the syntactic structure like phrase structure errors [26-29], and it is considered to reflect the brain’s response to the word category violations.

However, some authors have argued that such early ERP component may be an artifact [30]. As the onset of ELAN is quite early, some researchers have questioned whether it is related to syntactic processing given that even lexical access may take at least 200 ms [31-32]. However, it has also been shown that language comprehension system can predict the syntactic structure from some characteristics at an initial stage thus leading to very early syntactically-related activations [33]. These processing events are often referred to as basic syntactic processes starting with pre-processing of the syntactic structure, and it is often assumed that they necessarily precede semantic processing [34-35]. Finally, early first-pass syntactic parsing stage is believed to have a high degree of automaticity as it has been shown to be independent of focused attention on the input [27].

Similar early morphosyntactic effects have been found using the MMN response. This component, typically related to acoustic change detection and auditory short-term memory [36] is also known to reflect higher-level language processes [37-38]. For instance, the so-called *syntactic MMN* (sMMN) was found to be elicited by syntactic violations [28, 39-42]. Previous studies found that whenever the deviant sequence included a verb person/suffix agreement violation (e.g. ‘we *walks’), it caused a larger sMMN in comparison to the same verb presented in agreement with the pronoun (‘he walks’). The authors assumed that sMMN reflects the same early automatic processes as ELAN. Crucially, all of these effects were registered when the participants’ attention was diverted away from the auditory stimuli, typically using an active visual task. Furthermore, when directly comparing conditions with attention focused on vs. diverted away from the syntactic inputs, it was found that this early MMN/ELAN deflection was not affected by attention allocation until approximately 200 ms, implying a strong degree of automaticity in early syntactic parsing [42]. Such early automatic (morpho)syntactic activity may reflect processes associated with the listener’s attempts to maintain efficient alignment with the speaker and associated prediction (priming) mechanisms [43]. As such, they may have to rely upon activating syntactic templates and junctions as early as possible and often in a highly predictive and automatized fashion [40, 44].

Although such early automatic (morpho)syntactic processing has been repeatedly demonstrated in neurophysiological research, it remains unclear whether the listener’s brain makes use of extralinguistic speaker-related information during syntactic parsing in a similarly rapid automated fashion. Moreover, studies focused on extralinguistic processing are also limited with regard to the language in which the materials are presented. Languages with a shallow inflectional system (such as English, the most often used language in psycho- and neurolinguistic literature) lack important information about the processing of speaker-word gender incongruences. The present study is aimed to fill these gaps by examining the neurophysiological correlates of automatic extralinguistic processing using subject-verb agreement in Russian under non-attend design while carefully balancing acoustic, psycholinguistic and voice-related properties of the stimuli.

In particular, we investigated how and when linguistic and extralinguistic gender information, such as speaker-dependent voice congruency, can be integrated during sentence processing. For this purpose, we used auditory passive presentation protocol previously employed in sMMN and ELAN studies (see e.g. [28]). Participants were presented with short auditory phrases that may be syntactically felicitous or not while their attention was diverted away from the auditory stimuli to a primary visual task. Furthermore, by using male and female voices that either matched or mismatched the grammatical gender of the linguistic stimuli (e.g., ‘*I walked*_*MASC*_’ recorded in male or female voice), we directly assessed the integration of voice-related extralinguistic information into automatic morphosyntactic processes. Since gender-voice processing may occur quite early [13] supported by automatic morphosyntactic analysis, grammatical gender may also be identified in a similar manner – signaled by the verb’s morphology and in close integration with voice-gender information.

Our hypotheses were motivated by previous findings showing early automatic stages of morphosyntactic processing starting well before 200 ms [28, 39, 41, 45]. In case of bottom-up automatic processing, we expected to observe a similarly early automatic ELAN-like modulation in the voice-gender mismatch conditions. Alternatively, if the integration of voice information is carried out in a top-down manner, no effects should be expected at early stages of processing, and the speaker-related information could be assessed only after the syntactic structure analysis, either only modulating later responses in the N400-P600 range (similar to [17] study) or showing no modulation at all in our non-attend design.

## Method

### Participants

We recruited 37 right-handed participants (17 males, age range 19-32 years, mean age = 22.55, SD = 3.1) to take part in the experiment. All of them were Russian native speakers with no history of neurological or psychiatric disorders, normal or corrected-to-normal vision and normal hearing. All of them filled a consent form prior to the experiment. This research was approved by the HSE University Psychology Department Ethics Committee, Moscow, Russia.

### Materials

Ten Russian verbs comprised of 5 phonemes — *куnил* ([kʊp^j^ˈil], *bought), велел* ([v^j^ ˈl^j^el], *ordered*), *заnел* ([zɐˈp^j^el], *sang*), *надел* ([nɐˈd^j^el], *put on), nобил (*[pɐˈb^j^il], *broke*), *засел (*[zɐˈs^j^el], *sat*), *nоnал* ([pɐˈpal], *got*), *nолил* ([pɐˈl^j^il], *watered*), *nожал* ([pɐˈʐal], *shook*), *сумел* ([sʊˈm^j^el], *could*) — combined with first-person singular pronoun *я* ([ja], *I*) were selected as experimental materials (see S1 Appendix). In order to allow gender manipulation, all verbs were used in their past tense and consisted of a four-phoneme base and the past tense suffix *–л* ([-*l*]) with zero ending corresponding to the singular masculine gender form. All experimental stimuli were of identical duration, they were balanced by frequency, and they had high surface-form and bigram frequencies (see S1 Appendix). By combining the pronoun and the verbs (e.g., я куnил [ja kupil], *I bought*_*MASC*_) we constructed 2 sets of 10 sentences. All sentences were presented in two versions — recorded in both female and male voice (i.e., *I verb*_*MASC*_*_*by a male speaker and *I verb*_*MASC*_*_*by a female speaker) thus they only differed in the voice-gender agreement. In this instance, the verb’s morphology (suffix) provides information not only about the syntactic congruency but also about the speaker-gender one available by means of voice differentiation. Sentences were synthesized with the help of Voice Reader Home 15:Test voices (Linguatec) using male and female native Russian speakers. All phrases were grammatically correct in terms of their subject-verb agreement. However, given that there were two speaker’s genders – male and female, their grammatical form was either congruent or incongruent (equiprobably) to the gender of the speaker (see Fig 1 for experimental conditions and stimuli examples). Identical verbs appeared in congruent and incongruent conditions, differing only in the presentation voice.

**Fig 1.**
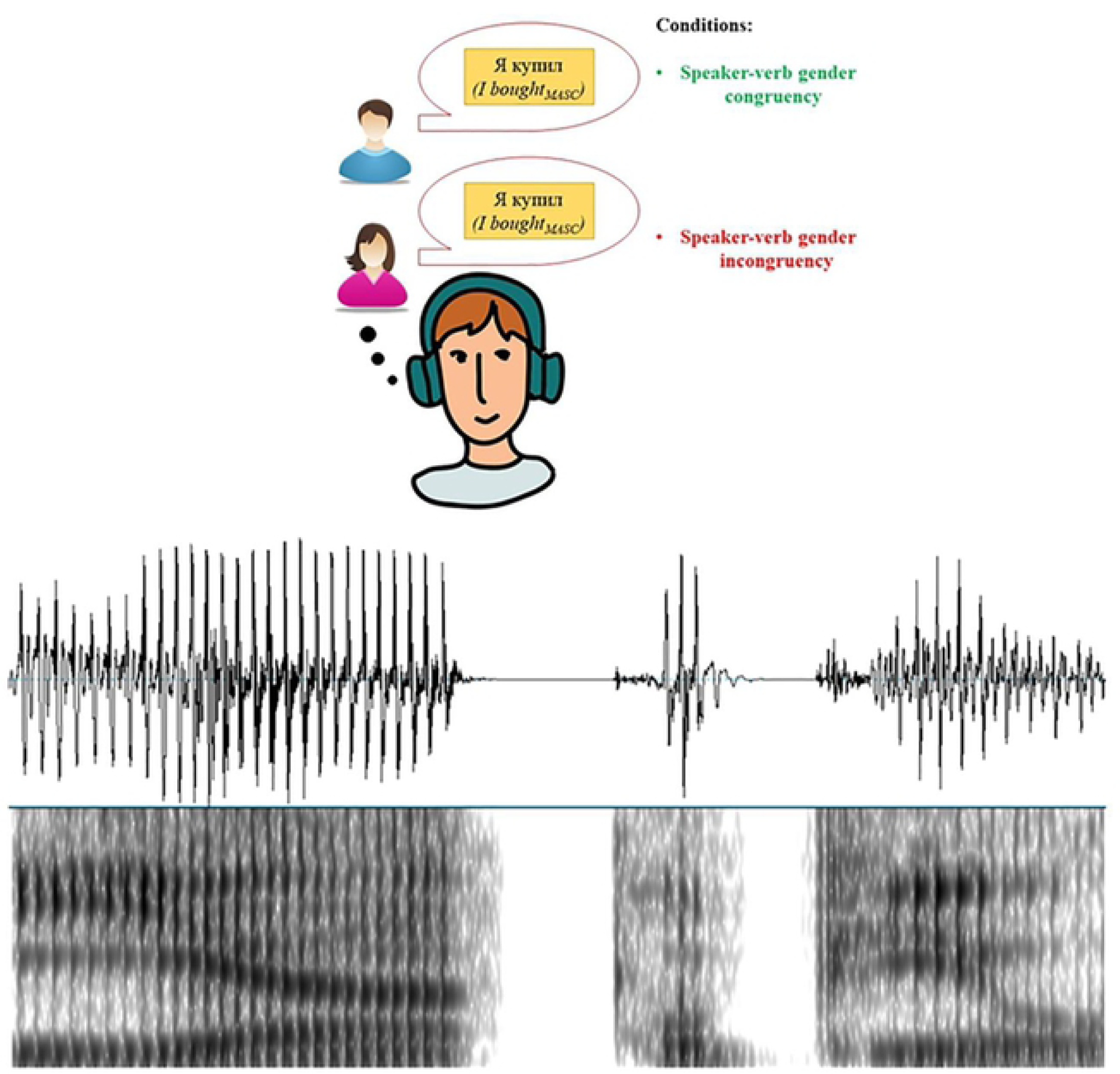
Example of experimental stimulus and its waveform and spectrogram (*я куnил* / *I bought*_*MASC*_) in male voice.

To minimize acoustic variability between stimuli, which could affect brain responses, pronouns and verbs were produced separately and then cross-spliced together into sentences by means of Audacity v2.3.0 software (Audacity Team). All the auditory stimuli were set in mono, .wav format. Stimulus duration was approximately 1000 ms (see S2 Appendix).

### Design and procedure

Participants were presented with the stimuli at a comfortable hearing level binaurally via earphones. They were tested in an electrically and acoustically insulated laboratory, and they were instructed to watch a silent film throughout the duration of the experiment (*Wallace and Gromit: The Curse of the Were-Rabbit)*, and to pay no attention to the auditory stimuli. The film was presented on a 28-inch Samsung-U28D590D LCD monitor with 1920 × 1080 resolution at 100 Hz refresh rate. Auditory stimuli were programmed and presented using E-prime v2.0 software (Psychology Software Tools, Inc., USA). During the entire experiment, the brain’s electrical activity was recorded using EEG (see details below).

We used 10 sentences presented in two voices during the experiment, each repeated 20 times in pseudorandom fashion, therefore 2*10*20 = 400 total trials. The average stimulus onset asynchrony (SOA) was 1050 ms, jittered to range from 1000 ms to 1100 ms. The sequence was subdivided into 4 blocks, 3.5 minutes each, to avoid participants’ fatigue. An initial training block with 10 trials was also carried out to familiarize participants with the paradigm consisting of similar verbs (but not identical to the experimental materials). A pseudo randomization of stimuli was established in each block such that the same phrase should not occur twice in a row.

There were self-regulated breaks between blocks, when participants were asked to fill a multiple (5) choice questionnaire about the content of the film. This ensured that participants indeed paid attention to the film and not to the auditory stimuli. The questionnaire contained a total of 20 questions with 5 questions per block. At the end of the task, participants were asked to carry out a word recognition task in order to provide a behavioral verification of the passive presentation of audio stimuli and to determinate that participants were sufficiently distracted from the auditory input. It consisted of 10 experimental verb forms plus 20 fillers (experimental verbs in another form and filler verbs). Participants had to read and indicate whether the item appeared in the experiment. The total duration of the experiment was ∼50 minutes excluding EEG set-up and preparation.

### EEG recordings and pre-processing

The brain’s activity was continuously recorded by means of 128 active electrodes, amplified using an actiCHamp EEG system (Brainproducts GmbH, Germany). Electrodes were mounted in a 128-channel cap (easyCap, Brainproducts GmbH, Germany) following the standard 10%-10% EEG configuration system. During the recording, EEG signals were sampled at 1000 Hz and a notch filter was applied at 50 Hz to remove line noise. Cz (vertex) was used as a reference electrode. Three electrodes were used for electrooculogram (EOG) recordings – two of them were placed on left and right canthi of the participants’ eyes for recording of horizontal ocular activity and another one under the participant’s right eye, for the recording of vertical ocular activity. Impedances were always kept below 10 kΩ. Stimulus markers were sent to the recording computer at the onset of the past tense verb suffix using E-prime software.

Preprocessing of the EEG data was conducted using BrainVision Analyser 2.1.2 software (Brain Products GmbH). First, high- and low-pass filters were applied at 0.1-30 Hz, following filtering parameters applied in similar studies [46-47]. New bipolar horizontal and vertical EOG signals were computed by subtracting differences between monopolar channels above (Right-Left for HEOG and VEOG-Fp2, respectively) using the Formula Evaluator function. Before artifact rejection, raw data were inspected with ± 100 µV amplitude and 100 µV of max-min difference as the artifact criteria within individual channels. This step ensured the detection of channels with sustained bad signal throughout recordings and their correction at a later step. Following raw data inspection, ocular correction ICA was used in order to remove all independent components reflecting ocular activity (saccades and blinks). Ocular channels or bad channels were not included into the ICA procedure. Then, triangular topographic interpolation was carried out to recover bad channels detected previously during raw data inspection. Afterwards, EEG data was epoched per participant and per condition (congruent and incongruent) using the 100 ms before and 800 ms relative to the onset of the past tense verb suffix. The 100 ms pre-suffix interval was used for baseline correction. Following this, artifact rejection was applied to each epoch, with the same criteria as for the raw data inspection. Then, an average reference was applied, with the EEG signal in each epoch re-referenced to the mean activity in all EEG channels. Finally, the remaining artifact-free epochs were averaged per each participant and condition in order to compute ERPs. The number of epochs used per each condition in each subject was not less than 180 out of 200.

### Data analysis

For unbiased data-driven analysis, overall activation was first quantified as the global field power (GFP) of the ERP responses across all scalp electrodes, stimuli, and conditions. To this end, the grand average response was first calculated across all conditions and stimuli collapsed and then the root mean square was calculated on the sum of squared amplitudes across all electrodes. Visual inspection of the GFP identified 2 distinct peaks – approximately at 150 ms and another approximately at 400 ms after the onset of the past tense verb suffix (see Fig 2). As note, the latency and topographic distribution of these effects corresponded with that found for both ELAN- and N400-like components in related ERP literature []. Thus, based on the inspection of the GFP waveform, we selected two time windows (130-170 ms and 350-450 ms) and extracted the mean activity from each participant, electrode, and condition and subjected them to further statistical analyses.

**Fig 2.**
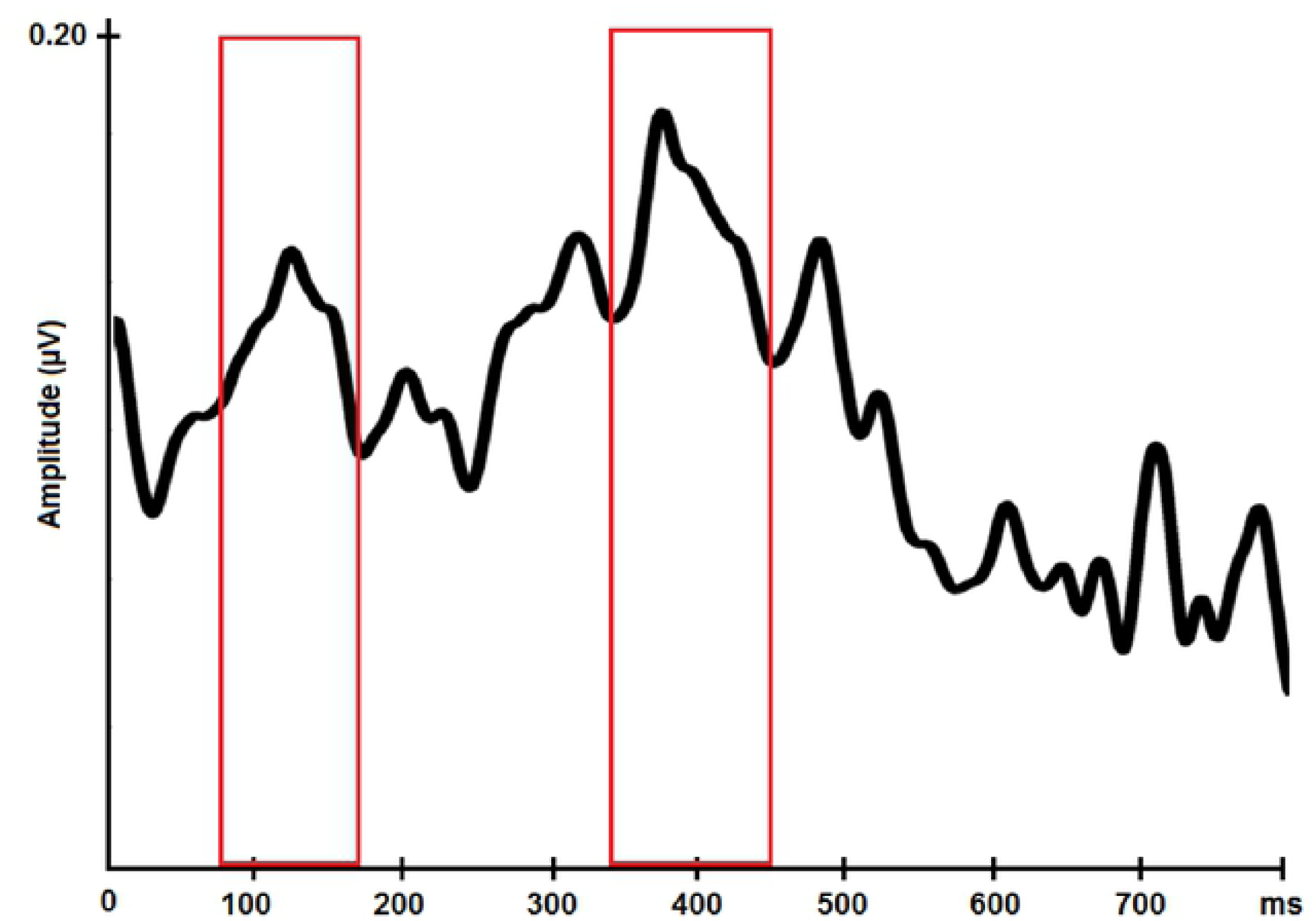
GFP waveform for the voice congruency effect. Red bars highlight two main peaks along the ERP segment, maximal at around 150 ms and 400 ms post-suffix onset.

Repeated-measures analyses of variance (rmANOVA) were carried out in order to analyze differences in mean activity between congruent and incongruent conditions, separately at each time window. To this end, window-mean amplitude data from an array of 64 electrodes were included into the statistical analysis, broadly distributed over frontal, central, and posterior areas where auditory ERPs are typically most pronounced. The selection of these regions for the analysis followed previous findings related to auditory syntactic processing. These electrodes were divided into eight topographical lines (F, FFC, FC, FCC, C, CCP, CP and CPP) with eight electrodes in each line. The mean activity for each electrode in each line was submitted to a 2 × 8 × 8 rmANOVA with factors Voice congruency (congruent vs. incongruent) × Anterior-Posterior (8 horizontal lines) × Laterality (8 electrodes - left to right), for the analysis of gender congruency depending on the speaker’s voice. All *p*-values were corrected for non-sphericity using Greenhouse-Geisser correction where appropriate. Effects reaching significance were followed-up with post-hoc-tests, employing Bonferroni correction for multiple comparisons.

## Results

### Behavioral Data

Behavioral data analyses included extracting mean scores from each participant in the film questionnaire and verb recognition task in order to assess their compliance with the task. Ratings obtained from film questionnaires indicated that participants were effectively paying attention to the video during auditory presentation (mean correct answers = 19.31 out of 20, SD = 0.3). Conversely, ratings obtained during the recognition task showed poor recognition of the auditory stimuli (experimental verb form: mean= 4.45 out of 10, SD = 1.3; experimental verb in another form: mean = 1.56 out of 5, SD = 1.2; filler verb: mean = 1.66 out of 15, SD = 1.5). Since all participants were paying attention to the film and not to the auditory stimulation data, all ERP data were used in further analysis.

## ERP data

## Results

### ELAN-like component (∼150 ms)

Fig 3 presents ERP waveforms and topographic distribution of brain responses at the early time window of 130-170 ms where an ELAN-like response was registered. rmANOVA indicated a significant three-way interaction between Voice Congruency, Anterior-Posterior and Laterality (*F*_49,1764_ = 2.634, *p* = .006, *η*2 = 0.068). Post-hoc comparisons indicated more negative responses for incongruent stimulus (*F*_1,36_= 5.620, *p* = .023, *η*2 = 0.135) at fronto-central, central, and centro-posterior regions in both hemispheres (FFC, FC, FCC, C, CCP, CP, CPP) with the strongest effect at central sites (see Fig 3A).

**Fig 3.**
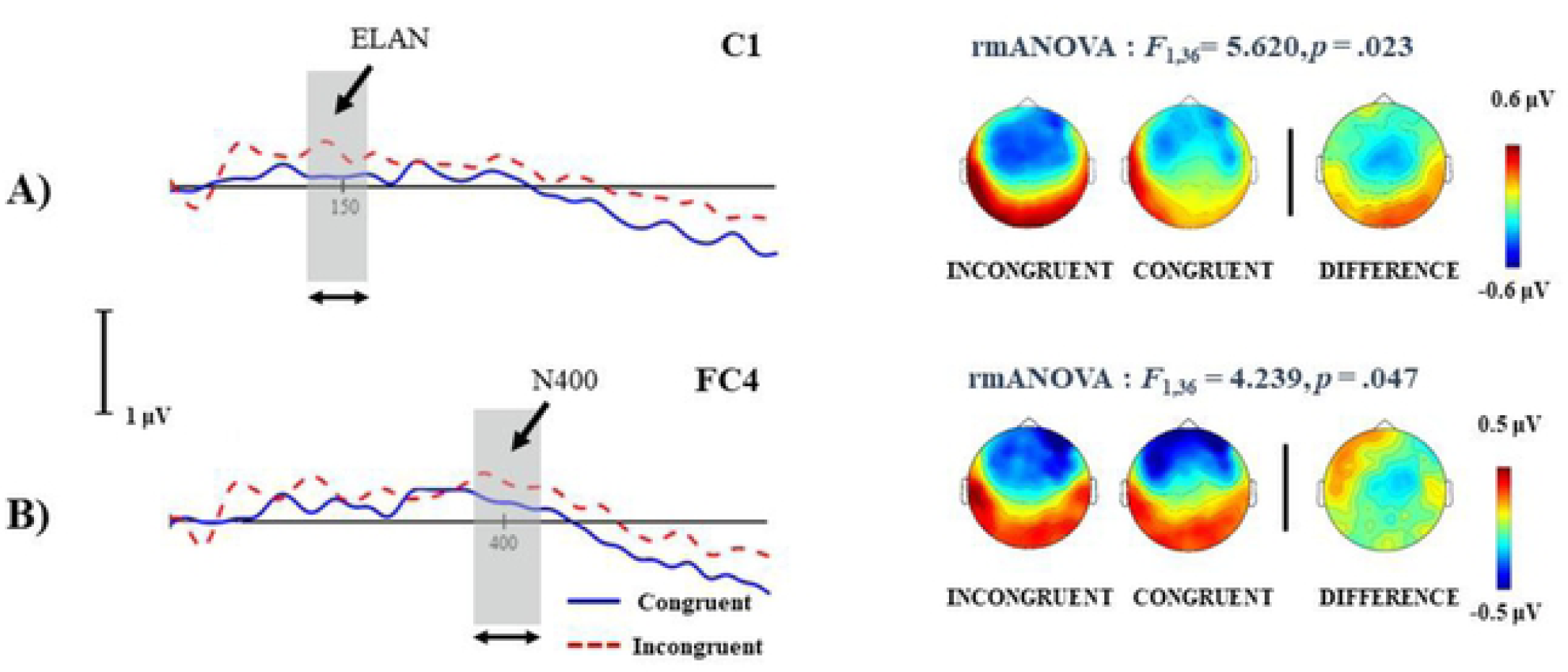
Average ERP waveforms and topographic maps for Congruent and Incongruent conditions. Grey shaded areas highlight early and late Voice Congruency effects at (A) 130-170 ms and (B) 350-450 ms after verb-suffix onset, compatible with ELAN and N400 components, respectively.

### N400 component (∼400 ms)

Analysis of the Voice Congruency effect at the later time window (350-450 ms) revealed in a significant three-way interaction between Voice Congruency, Anterior-Posterior and Laterality (*F*_49,1764_ = 2.072, *p =* .024, *η*^2^ = 0.054). Post-hoc comparisons showed stronger negative responses for stimulus in disagreement with speaker’s gender (*F*_1,36_ = 4.239, *p =* .047, η^2^ = 0.105) at right fronto-central and central regions (FC, FCC, C) with particularly strong effect at right central sites (see Fig 3B).

## Discussion

The current study used Russian – a morphologically explicit language with a rich gender-marking system – in order to investigate online neural dynamics elicited by automatic syntactic processing under the influence of extralinguistic gender information provided by the speaker’s voice. Using passive auditory presentation protocol and a set of controlled first-person pronoun-verb phrases, where the verb gender marking was either congruent or incongruent with the speaker’s voice, we investigated the putative speaker-voice congruency effect in the brain’s ERP responses to spoken stimuli. Analysis of the speaker-voice congruency contrast revealed both early and late ERP effects indicating increased negativity in the voice-gender incongruent conditions. We will briefly summarize and discuss the main findings of the study below.

A significant incongruency effect was found as earlier as 130-150 ms after recognition of grammatical gender, likely lead by the identification of verb’s morphology (suffix onset). Such latency broadly corresponds to the time window of the well-known ELAN component, which typically shows an enhancement of the negativity for syntactically incongruent vs. congruent stimuli over anterior channels, as it was also the case in our study. To the best of our knowledge, this is the first demonstration of this ERP effect for *extralinguistic information*, with previous findings revealing only later effects during syntactic processing [14-17]. At the same time, this finding is in line with previous studies reporting ELAN/sMMN effects for agreement violations [28, 38-39, 42, 48-49]; however, in the present study the early agreement effect was elicited by voice incongruencies in otherwise grammatically sound phrases and not by a morphosyntactically infelicitious agreeement.

A mechanistic explanation of early syntactic responses has been offered by the so-called *(morpho)syntactic priming* hypothesis [39], claiming that previous information (e.g., subject) leads to pre-activation of the relevant affix representation in order to facilitate and expedite input processing. Thus, when the affix finally arrives, it has been in part preactivated and less activation is necessary relative to pre-affix baseline. When such a pre-activation is not possible, the representation of the unprimed affix (or any other unexpected morpheme) has to be activated from a lower functional state manifesting as a larger activation relatively to the pre-affix baseline. Thus, a relatively larger response is registered for non-preactivated items in incongruent combinations. Whereas this original proposal was based on the existence of associative links between related morpheme representations (formed through their co-activation during previous language experience; e.g., singular pronoun and singular verb affixes in pronoun-verb phrases), it can be expanded to extralinguistic information. In this sense, we may hypothesize that a female/male voice pre-activates feminine/masculine-gender morphology, respectively. As a result, a prediction is made: The speaker’s gender is used to automatically predict the upcoming verb’s morphology in order to maintain interlocutional alignment [43]. When this prediction is not supported by the auditory input, an ELAN-like response is observed similar to that found for syntactic violations. Such an early chronometry of this response supports the notion of activating syntactic prediction templates (syntactic representation of a particular grammatical structure) which are available to the processor even before the input unequivocally supports this prediction [44].

Other aspects of the speaker-dependent information may be also pre-processed as particular language templates, and their violation should be accompanied by similarly early electrophysiological responses as suggested by some previous MMN studies (e.g., [13]). Note, however, that a very early onset of the observed modulation could also reflect a high predictability of the stimuli with the voice gender identity being clear from the pronoun onset and before the gender-specific ending onset. Whereas this per se does not undermine the result, which can only suggest that this is what happens in real conversation settings, further studies are necessary to ascertain this and could use a larger choice of forms using different genders, as well as possibly different voices in both congruent and incongruent conditions with more stimulus variability.

Another important aspect of our findings is the fact that the present early ELAN-like response was found in the outside of attentional focus on the linguistic input as the participants were engaged with the primary visual task. This result reinforces the findings of other studies that documented grammatical morphosyntactic effects for unattended agreement violations [40, 42, 45]. One important difference, however, is that the effect reported in this paper was elicited by extralinguistic rather than purely morphosyntactic factors. The early time-course of the effect and its emergence outside of the attentional focus indicate a high degree of automaticity of extralinguistic feature’s integration into sentence processing.

We have also registered a congruency effect at a later time window (350-450 ms) as a more negative response for voice incongruent stimuli than for congruent ones at later stages of the processing. This effect, compatible with the N400 frame, is similar to that previously observed for voice-gender agreement [17]. In line with previous studies, this N400-like effect may reflect the integration of extralinguistic information at a secondary stage of sentence processing [16-17]. Although N400-like effects are typically distributed at parietal scalp sites, the modulation reported here showed fronto-central activation with more right-hemispheric topography. Such right-hemispheric distribution may be related by the well-known right hemispheric laterality of the voice and prosody processing [50]. In terms of more frontal activation, this negativity might result from action processing (verbal phrases), which is known to lead more anterior N400 effects [51]. Moreover, relatively more frontal N400-like effects have been previously reported for auditory stimulus presentation, such as implemented in the current study, also leading to longer latencies of the component in auditory domain [52-54]. Importantly, previous studies reporting N400 effects for extralinguistic integration found these responses during active linguistic processing (auditory attended presentation task) while the current study implemented passive presentation.

Overall, co-registration of both early ELAN-like and later N400-like effects suggests that processes underlying the integration of speaker-dependent features into sentence processing may be a two-stage *detection+integration process* commencing at early automatic detection stages and with the later deflection reflecting a second integration stage of this same process. Importantly, later effects are often considered as reflecting more controlled processing (see, e.g., [27, 42], for automatic and controlled syntactic ERPs) albeit they can be also generated under a passive demands-free task. To fully attest the automaticity of these processes, future studies will need to implement an explicit attention manipulation and directly compare the brain’s responses to attended and unattended voice-syntax (in)congruences.

## Conclusion

The present ERP study provides electrophysiological evidence showing how extralinguistic information is integrated into the neural processing of syntactic information by the human brain. ERP analysis of brain responses to voice-congruent and voice-incongruent pronoun-verb phrases revealed two distinct processing stages: An early automatic stage (reflected at the ELAN time window) and a later stage (in an N400 time window), both showing enhanced frontal negativity for the mismatching stimuli. These findings (1) confirm previous results and, (2) provide novel evidence regarding the neurophysiological bases of syntactic processing. Importantly, we show that the latter does not only rely on the available linguistic information but also on the presence of extralinguistic cues; such as, the speaker’s gender information assumed from their voice. Moreover, this integration appears to take place in a rapid and automatic fashion as evident from the timing of the effects and their presence in the absence of focused attention on the stimuli. Nevertheless, further investigations are necessary in order to better understand the neural bases of this interaction. The use of more complex paradigms, including more variable morphosyntactic contrasts, variability in voices and as well as brain imaging techniques with high resolution at both temporal and spatial dimensions (such as MEG with MR-based source reconstruction) will lead to a better understanding of spoken language comprehension in the human brain.

## Acknowledgments

Supported by the HSE Basic Research Program and the Russian Academic Excellence Project ‘5-100’.

## Supporting information

**S1 Appendix A. Full set of words used in stimuli**.

**S2 Appendix. Full set of stimuli phrases with duration**.

